# Neighbor sensing through rhizodeposits in sorghum affects plant physiology and productivity

**DOI:** 10.1101/2024.09.04.611020

**Authors:** Shiran Ben-Zeev, Amanda Penn, Erica H. Lawrence-Paul, Desa Rae Abrams, Rotem Ben-Zeev, Carolyn Lowry, Jesse R. Lasky

## Abstract

Plant-plant interactions play a crucial role in shaping the growth environment for crops, impacting their productivity and stress response. The interaction between plants aboveground has been studied and incorporated into breeding programs that select for plants that reduce aboveground competition between plants. However, few studies have focused on belowground interactions, and these looked at combined interactions and root partitioning in the soil. This study focuses on the developmental and physiological responses of sorghum (*Sorghum bicolor* L.) genotypes to neighboring sorghum plants. In this study, we used two growing methods: i) a focal plant surrounded by neighboring plants in the same pot but without shading, and ii) a focal plant grown either alone or surrounded by neighbors, irrigated with nutrient solution passed through pots (leachates) with or without plants. Our results showed that the presence of neighbors in the same pot led to reduced size-dry weight height, and leaf area of the focal plant. In addition, the presence of neighbors reduced stomatal conductance and PSII quantum yield. While the response direction was similar across tested genotypes, the magnitude varied. The results were repeated when neighboring plants were not grown in the same pot, but the nutrient solution passed through the root system of plants of the neighboring genotype. Furthermore, we saw a reduction in assimilation rate and stomatal conductance when plants were exposed to either the physical presence of neighbors or leachate. We did not find differences in root architecture in either treatment. These results show plants change their growth in response to neighbors and that the signal is carried through the liquid phase of the soil. Our findings provide insights into sorghum plants’ responses to below-ground signaling from neighboring plants and lay the foundation for future studies enabling increased crop performance under high-density planting conditions.

## Introduction

The interactions between a plant and its neighbors define much of the immediate environment in which it grows. This environment can be beneficial or disadvantageous, depending on the identity of the surrounding plants and the plant’s ability to cope with the constraints of competition or utilize the benefits of facilitation (Farrior 2014). Global climate change and the need to produce more food from less land require increased farming intensity. One approach to improve productivity has been developing crops that yield more at the stand level when grown in high densities, even if individual plants yield less. Since the 1960s, ideotypes-putatively ideal combinations of crop phenotypes - have been developed to reduce competition and allow the growth of more plants per unit area (Donald 1968). Plants compete for sunlight above ground by increasing leaf area and stem height. Thus, many traits have been targeted in breeding programs, such as lower leaf angles to avoid mutual shading and smaller tassels in maize (Sandhu and Dhillon 2021). Understanding the developmental and physiological response to neighbors and its underlying genetic basis could enable further breeding for higher performance under intense competition for future farming.

Competition between plants, by definition, leads to reductions in fitness (Casper and Jackson 1997). Competition may become more severe when such resources are limited in supply and availability (Callaway and Walker 1997). Plant-plant competition can affect stand-level productivity by individual plants inhibiting the growth of plants around them through increased uptake of nutrients or water, causing scarcity, reducing stand-level productivity, or exuding allelopathic chemicals affecting the germination, growth, or development of neighboring plants (Putnam 1985; Bogatek *et al*. 2006). An opposite effect is facilitation among neighbors, e.g. by recruiting beneficial microbes, changing the pH, or changing soil structure in a way that benefits all plants in a stand (Farrior 2014; Henriksson *et al*. 2021). A recent study (Lorts and Lasky 2020) demonstrated that an *Arabidopsis* plant’s response to neighboring grasses leads to changes in aboveground and belowground organs, producing shorter inflorescence and reducing root diameter. The same study also showed that this response changes under abiotic stress, such as drought.

While resource competition has been the primary focus of ideotype development, belowground signaling among neighboring plants may also influence productivity (Bhowmik and Doll 1984) but is relatively less studied. Belowground resource competition mechanisms include increasing root system size and reducing the size or changing the direction of competitors’ root systems. The latter are changes that can play a vital role under a combination of competition and drought stress (Craine and Dybzinski 2013). Plants often respond to the presence of neighbors by altering their growth. A well-known response that affects both root and shoot allocation is shade avoidance (Page *et al*. 2011). In response to higher far-red red light, young maize plants developed a higher root-to-shoot ratio and produced a higher leaf area. Several studies have addressed changes in root gene expression, metabolites, and proteins in the presence of neighbors (Schenk 2006; Afifi and Swanton 2011; Khashi U Rahman *et al*. 2019). Schmid *et al*. (2013) presented results from Arabidopsis plants, which responded to the presence of a neighbor plant in the same pot by up or down-regulating genes involved in response to parasitic fungi, salt, heat, hypoxia, and phosphorus deficiency. The same study also presented a decrease in shoot biomass and changes to root morphology in Arabidopsis plants interacting with roots of *Hieracium pilosella*.

Responses to neighbors start at the seed stage (Hilhorst and Karssen 2000), leading to growth changes in seedlings, suggesting that root recognition precedes the aboveground encounters (Afifi and Swanton 2011) and sensing of neighbors (Schmid *et al*. 2013; Bilas *et al*. 2021). The presence of roots belonging to a different species drives changes in root distribution to different depths, similar to the changes caused by nutrient and water availability and placement (Mahall and Callaway 1991). In the rhizosphere, sensing of roots occurs via signals such as nutrient deficiency, water scarcity, and, importantly, chemical sensing of small recognition molecules and specific proteins secreted in root exudates (Badri *et al*. 2012). Lectins (e.g., jacalin), peroxidases, and jasmonic acid have been suggested as molecules mediating root responses to neighbors (including weed species in fields) (Bilas *et al*. 2021). Another important set of molecules affecting neighboring plants’ growth regulation are strigolactones (SL)(Proust *et al*. 2011; Al-Babili and Bouwmeester 2015). SLs are a class of plant hormones that have been shown to mediate interspecies plant-plant communication (Wheeldon *et al*. 2022; Yoneyama *et al*. 2022) and parasite-host recognition (Babiker and Hamdoun 1983; Bellis *et al*. 2020; Mwangangi *et al*. 2023) and are involved in plant-plant interaction and development in the presence of neighbors (Bilas *et al*. 2021). The effect of root-root communication on flowering times has been reported in *Brassica napa* (L.), such that day-length sensing could be mimicked when exudates from plants sensing a long day are used to irrigate plants grown under short days leading to significantly earlier bolting and flowering (Falik *et al*. 2014).

Sorghum (*Sorghum bicolor* L. (Monech)) is a C4 cereal grown in rain-fed and irrigated environments on high-yielding and marginal soils from Africa and the Middle East to India, Australia and America. It is the fifth most economically important crop in the world (FAOSTAT, https://www.fao.org/faostat/en/#data), extensively grown in various sectors such as biofuel production, forage cultivation, and grain production. Domesticated in Africa (De Wet and Harlan 1971), where it is still grown on vast areas of land, sorghum is more resilient to environmental stress than many other crops, making it an important model species for studying the unique strategies for dealing with the drought stress of C4 grasses (Pardo and VanBuren 2021). Studies on sowing densities in sorghum, which are natural test cases for plant-plant interactions, have shown that sorghum harvest indices reduce when water is limited and that this reduction is uniform across sowing densities (Berenguer and Faci 2001), suggesting an individual-level response to biotic stress. However, high sowing densities have also been shown to result in yield reduction across three drought treatments (Jahanzad *et al*. 2013), representing a constraint of more neighbors. Two other studies reported a lack of response or increased productivity under higher densities (Gerik and Neely 1987; Raphaël *et al*. 2024), indicating a variability in the response to density.

This study was conducted to assess the effects of drought and perceived competition on plant growth across genotypes. We hypothesized that sorghum has various pathways to sense and respond to changes in its biotic environment above and below ground. To study these responses, we tested three subsequent hypotheses:

i. Sorghum plants will grow taller in response to the presence of neighbors, which would promote light capture even without sensing shading.
ii. Sensing neighboring plants below ground will serve as a sufficient signal to lead to changes in root and shoot growth, plant physiology, and reduced productivity.
iii. The effects of neighboring plants will be larger under water limitation.
iv. Biochemical signals present in leachates can trigger physiological and developmental responses to neighbors.

Our goal in these experiments was to observe the growth, development, architecture, and physiology of sorghum plants in systems where the focal plants were allowed to communicate and interact with their neighbors through signaling in the rhizosphere while preventing above-ground shading (by forcing neighbor plants away from the tested focal plant) and creating a uniform environment of above ground volatile compounds (through a random block design). Further, we sought to isolate the effects of sensing rhizodeposits, root exudates and other bio-chemicals released into the soil by a plant (Duchene *et al*. 2017). To achieve that, we studied plants without physical neighbors in the same pot. We accomplished this by transferring leachates from source pots with “neighbor” plants to pots of focal plants with and without the physical presence of neighbors (Fig S1).

## Materials and Methods

### EXPERIMENT 1- DROUGHT & PHYSICAL INTERACTION (EXP1)

#### Plant material

Ten sorghum genotypes obtained from USDA GRIN were selected to represent a wide range of variation. They included genome-sequenced grain sorghum lines RTx430 and BTx623 (PI 655996, and 564163, respectively), well-studied lines Shanqui Red (PI 656025) and SRN-39 (PI 656027), and landraces from India, Sudan, Nigeria, and South Africa (Table S1). These lines are referred to hereafter as focal plants for which data was collected. In addition, one common sorghum line (BTx 3440), received from William L. Rooney at Texas A&M, was used as the “neighbor” line across all experiments to avoid variation in signals and growth habit. The competitor line was selected for its high performance under drought conditions and seed availability.

#### Experimental design and growing conditions

Our goal with this experiment was to grow sorghum plants of a focal genotype with neighboring plants in the same pot, allowing them to interact below ground but preventing mutual shading and competition for light. Further, we tested the effect of drought on competition by including well-watered and water-limited treatments.

The experiment included four treatments in a full factorial randomized block design, including four blocks with one repetition of each genotype X treatment included in each randomized block. The treatments were: well-watered without competition (NCW), well-watered with competition (CW), drought without competition (NCD), and drought with competition (CD).

The competition treatments were created by surrounding a focal genotype with five neighbor plants (equivalent to 16 plants per square meter plant density or approximately 125,000 plants per ha), with the recommended range for sorghum sowing in the USA being 125,000-200,000 plants/ha in well-watered environments (https://extension.psu.edu/forage-sorghum-planting) (Figure S1). To avoid the effects of shading by the neighboring plants and focus our experiments on belowground interactions, the shoots of neighboring plants were trained to grow away from the focal plant by placing a paperboard bowl with a hole cut in its middle point so that the focal plant grows through the bowl and competitor plants are forced to grow sideways. At a later stage of growing (∼25 days after sowing), neighbor plants were restrained toward the outside of the pot using string to prevent them from shading the focal plant.

Seeds were sown in vermiculite in bottom-watered pots. Five days after sowing, seedlings were transplanted into 5.7 L pots (Fig S1 left) filled with 80% potting mix (Pro-mix BX, PA, USA) and 20% Turface© (Turface Athletics, IL, USA). To avoid any nutrient limitation, the soil in each pot was mixed with 14g of Osmocote © flower and vegetable 14-14-14 slow-release fertilizer (ICL, Israel), a high fertilizer rate selected to lead to high nutrient availability for the entire growth period. The dry weight was measured for each pot, and the volume required for saturation was tested and found to be 89% of the dry weight. At the start of the experiment, pots in the well-watered treatment were watered with the volume required for saturation. For the drought treatment, 75% of the saturation volume of water was added to allow seedlings to establish without saturating the soil. Pots were weighed every 3-4 days.

Drought treatment pots were maintained at 25-35% of the weight at saturation (once they have dried sufficiently (17 days after planting), and well-watered pots were maintained at 55-70%. The water deficit was validated using three soil volumetric moisture probes for each treatment (ZL6 Basic, Meter Environment, WA, USA), confirming the weight measurements.

Plants were grown in a greenhouse with day/night temperatures set to 28/18°C, with added light (3000 lumens) supplied by HPS lights to produce a 14-hour day length for 35 days after sowing in each cycle; the first cycle started on 27 December 2022, and the last cycle was harvested on 23 May 2023.

#### Phenotype measurements

For the focal plant in each pot, height was recorded as the distance from the soil to the base of the top fully expanded leaf. Height was measured every 3-4 days to calculate the height growth rate. Stomatal conductance and chlorophyll fluorescence were measured under full light (9-10 am) using a Li-Cor 600 fluorometer porometer (Li-Cor, Lincoln, NE, USA) as soon as leaves were wide enough to cover the sensor (32 days after sowing). At harvest, the height was recorded, and leaf area was measured. For EXP 1, leaf area was determined using a Li-Cor 2000 Plant Canopy Analyzer (Li-Cor, Lincoln, NE, USA). Shoot dry weight was measured using a bench top scale after drying the plants in an oven at 60 ° C for 3 days.

### EXPERIMENT 2- LEACHATE TRANSFER (EXP2)

#### Plant material

Two of the lines used in EXP1 were selected - RTx430 and BTx623 (PI 655996 and 564163, respectively), along with the recurring neighboring genotype, BTx 3440.

#### Experimental design and growing conditions

Following the results from the physical competition experiment, we designed a second experiment to separate neighboring plants’ physical and biological-bio-chemical presence on focal plants. This was achieved by transferring the leachate, likely including rhizodeposits, from a source pot of neighbors to a target pot with a focal plant. The system included two sets of pots: source pots positioned above drain holes (funnels) on 30 cm high raised benches and target pots positioned next to source pots on the growth table below. Leachates from source pots flowed into funnels and through a PVC tube to the target pots (Figure S1 right). The irrigation solution was transferred through media-filled pots with no plants for the control treatments and pots with three Btx3440 “neighbor” plants for the leachate treatment. To allow root visualization, the growth media was 60% sand (pool filter sand, Quikrete, Atlanta, GA, USA) 20% Turface© (Turface Athletics, IL, USA), and 20% clay (Kaolin calcinated clay, CB minerals LLC, Mamaroneck, NY, USA). Pots were determined to require ∼150 ml of water to be saturated, and thus, source pots were irrigated with 300 ml nutrient solution (Miracle-gro Aero Garden, Liquid Plant Food-Miracle-Gro, Marysville, OH, at 2ml/14L) every two days to avoid water and nutrient stress. Water amounts passing through the source pots were measured weekly to validate that there were no differences in the amount of water applied to experimental pots between control and leachate treatments, and the percentage of water passing was consistently 65-75% of the applied water. Target pots were only irrigated through source pots. Plants were harvested at 37 days after sowing.

#### Phenotype measurements

For the focal plant in each pot, the same phenotypes as EXP1 were collected. In addition, in-depth gas exchange measurements were collected on new fully expanded leaves using a LI-6800 Portable Photosynthesis System gas exchange (Li-Cor, Lincoln, NE, USA). These measurements included carbon assimilation rate, stomatal conductance, and transpiration rate for three plants for every treatment by genotype combination, as well as instantaneous water use efficiency calculated by dividing the assimilation rate by the transpiration rate (Farquhar and Richards 1984).)

At harvest (37 DAS), plants were removed intact from their containers, and roots were washed to remove soil. The roots of plants of the leachate transfer experiment (5 reps) were placed into sheet protectors (transparent bags) and scanned at 600 dpi, using the Epson 600V scanner (Seiko Epson Corp., Nagano, Japan). Images were analyzed using RhizoVision Explorer v2.0.3 for total root area, median root diameter, and number of root tips (Seethepalli and York 2020) using algorithms described by (Seethepalli *et al*. 2021).

#### Statistical analysis

Mixed linear models were used to estimate the effects of competition and drought; for EXP1, the model included genotype, competition, and drought as fixed effects, along with their interaction, with blocks as a random effect. For EXP2, the model included treatment and genotype as fixed effects along with their interaction, with block as a random effect. We calculated the LS means values using a model that included genotype, drought, and competition, as well as their interaction as fixed effects and the experimental blocks as a random effect. Models and hypothesis testing using the Tukey HSD tests and calculating LS means was conducted using JMP ® (SAS Institute, Cary, NC, USA,1989–2024). We compared the different models used for LSmeans using the R software (R Core Team 2021) lmertest package (Kuznetsova *et al*. 2015). We then used the ggplot2 (Wickham *et al*. 2016) package for plotting and significance was calculated with the ggsignif (Ahlmann-Eltze and Patil 2021) package. Correlations were calculated using the Corrplot package in R (Wei *et al*. 2017). The lsmeans package (Lenth and Lenth 2018) was used to calculate the cumulative fixed effects values for reaction norms analysis.

## Results

### EXPERIMENT 1

#### Neighboring plants reduce performance equivalent to a water-limited treatment

Competition in well-watered conditions (WW_C) significantly reduced shoot dry weight (F ratio= 45.3, P value = <0.0001) compared to the no competition-well-watered treatment (WW_NC), despite the ample nutrient fertilization provided in the competition treatment pots. Dry weight was reduced in both water-limited (WL) treatments compared to WW treatments, with no significant difference between the WL competition (C) and NC (Fig 1A, Table S2). For dry weight, these values were 73%, 59%, and 43% lower than the control in the WL C, WL NC, and WW C treatments, respectively. Plant height at the end of the growth period (34 days after sowing -DAS) was similarly reduced in response to competition and water limitation with a 20% reduction in WW C and 25% reduction in WW C treatments compared to control (F= 62.1 P<0.0001). A further significant reduction was observed under the combined competition and drought-WL C (40%) treatment (Fig 1B, Table S2).

**Figure 1:**
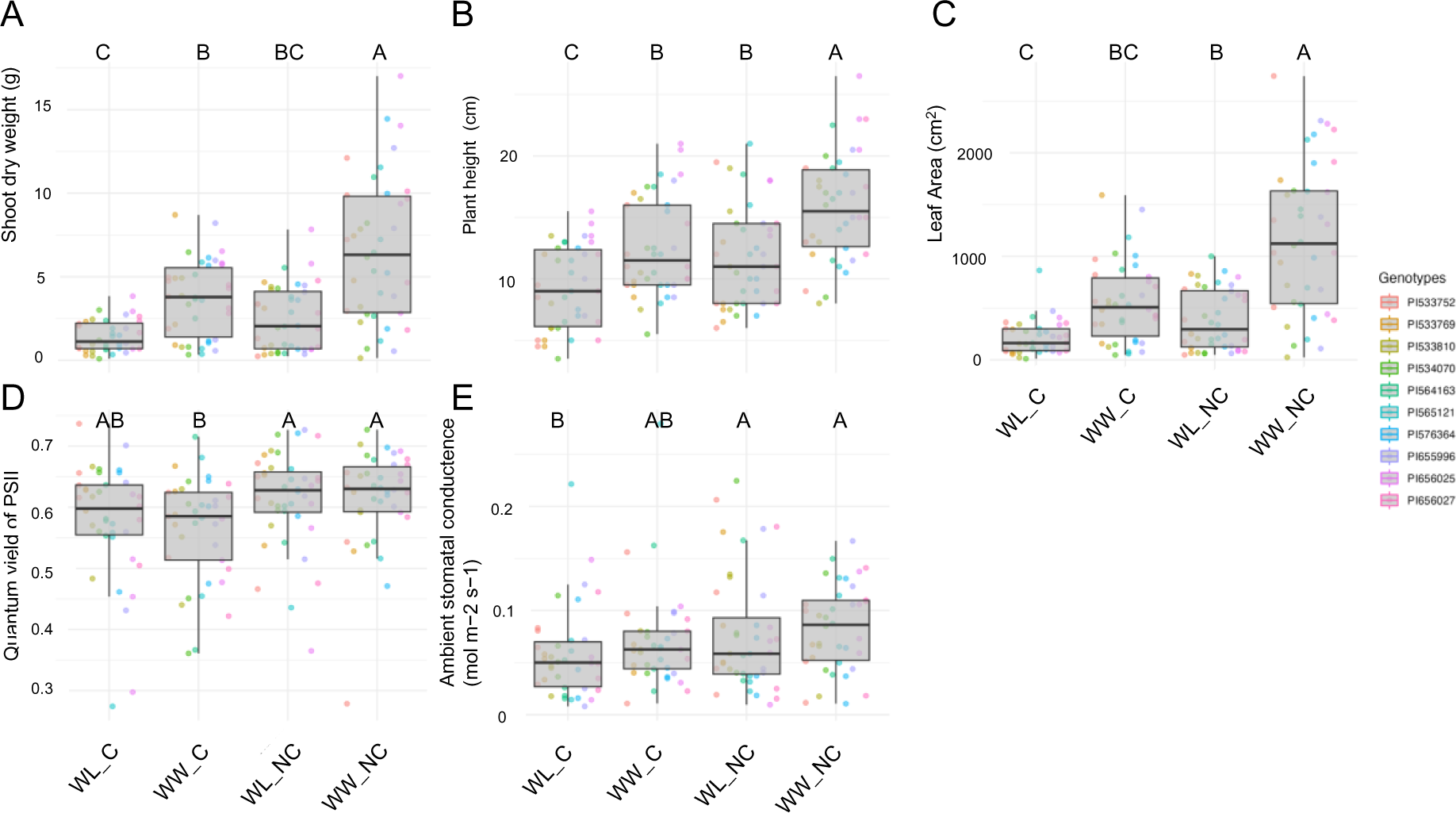
A—Shoot Dry weight (g), B—Plant Height (cm), C—Leaf area, D— Quantum yield of photosystem 2, and E—Ambient stomatal conductance, all measured 34 Days After Sowing (DAS). Values presented are all four reps of ten genotypes under water-limited (WL) and well-watered (WW) conditions with competitor plants (Competition) and without (No Competition). Connecting letters represent significance in Tukey HSD tests between treatments (P values lower than 0.05).

Leaf area was significantly lower in all treatments compared to the WW NC control, with the WL NC treatment being significantly lower than the control (65% reduction), the WL C treatment being significantly lower than both WW (82% reduction) and WL NC (F=64.4, P<0.0001). The WW C treatment (55% reduction) was in between the WL treatments and significantly lower than WW NC (Fig 1C, Table S2).

Similar to dry weight, competition led to a significant decrease in the quantum yield of photosystem II (PhiPSII) (Fig 1D, Table S2) under the well-watered treatment (representing an LS means difference of 8%) but not under the WL treatment (F=10.31, P=0.0016). Stomatal conductance was lower under the competition treatment in both water treatments. A 25% reduction was found for the WL C, respectively, compared to the control (F=6.31, P=0.01-Fig 1E, Table S2).

#### Variation in competition response among genotypes and between water treatments

We included genotypes from different regions and farming systems to test genotypic differences. However, the genotype effect was only significant for plant height, and none of the genotype-by-treatment interactions were significant, indicating similar responses to neighbors. This overall trend was generally uniform in direction (no significant genotype x competition or genotype x drought interaction found-data not shown). However, some differences were observed between genotypes in the size of the effects. For example, dry weight at 34 DAS was lower in the competition treatment under both water treatments, with a uniform response for all lines under the WW condition and more variable under WL (Fig 2). PI533810, PI533769, and PI534070 had a large difference in response to competition in the WW treatment and almost no difference under WL (Figures 2A and 2B). Plant height was lower in the competition treatment for all lines under both water treatments; PI576364 and PI533769 had smaller responses to the presence of neighbors but did respond to the water treatment (Fig 2 C, D).

**Figure 2:**
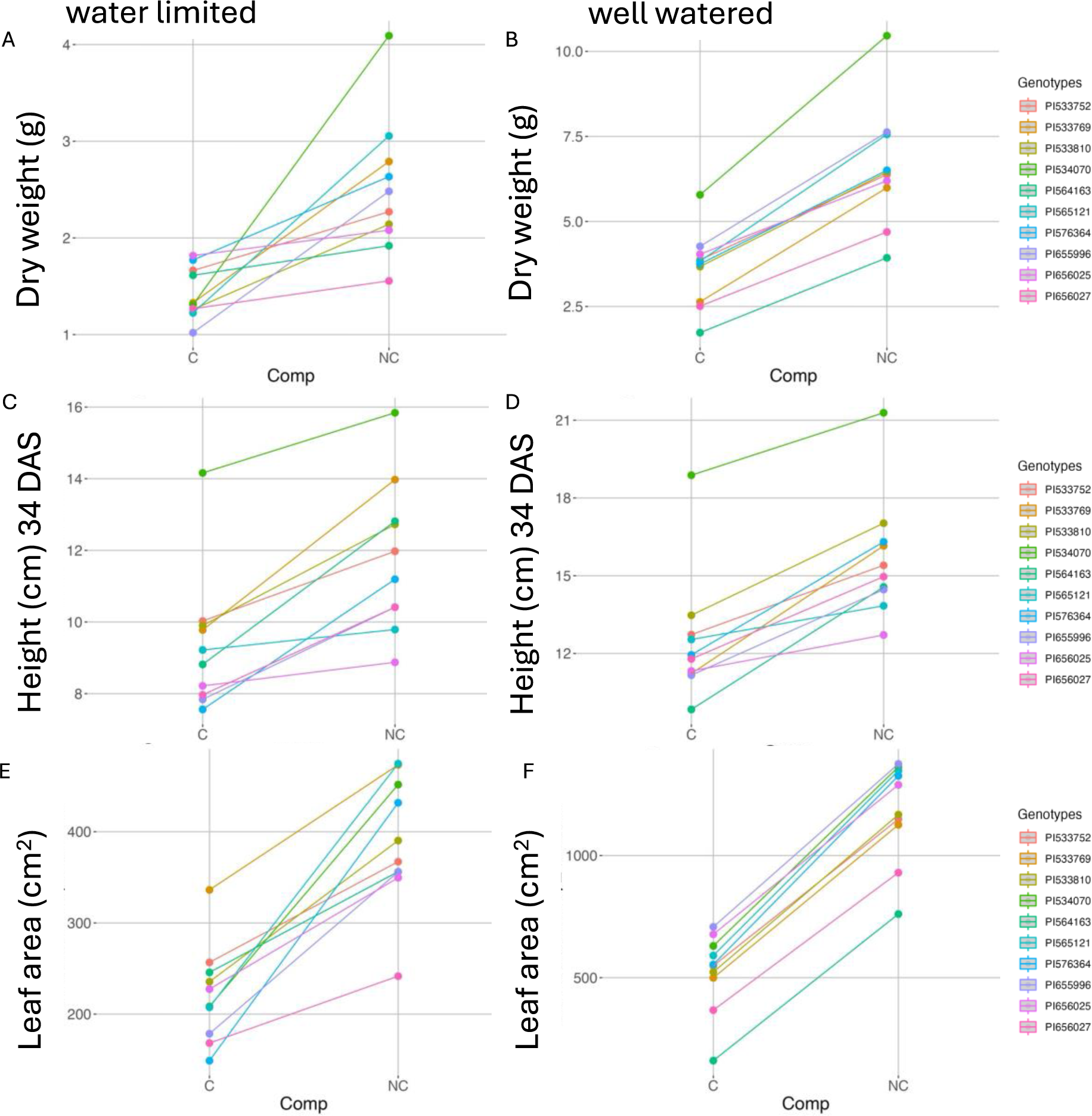
Reaction norms of the ten tested genotypes for dry weight (A, B), height (C, D), and leaf area (E, F) between the competition (C) and no competition (NC) treatment within the water-limited (WL, left), well-watered (WW, right) treatments. Values presented for each line are the least square means obtained from a model, which included the lines (PI numbers for each line are presented in Table S1), competition, experiment, water treatment factors, and competition X genotype interaction.

Leaf area was lower in the competition treatment under WL and WW (Fig 2 E, F). The response to competition under WL was variable between lines, with lines PI656025, PI576364, and PI 656027 having a large response and PI534070 having the smallest response. Under WW, the response to neighboring plants was mostly uniform across all lines.

### EXPERIMENT 2

#### Irrigation with leachates leads to a response to neighbors without plants growing in the same pot

Based on the reduction in plant productivity in response to the presence of neighboring plants, even in treatments with ample resources and in the absence of shading, we hypothesized that the signal for the presence of neighbors could be mimicked by transferring root rhizodeposits to pots with individual plants growing in them. Shoot dry weight was significantly (P<0.05) reduced by irrigating with leachate (Fig 3 A, Table S2) (F ratio = 11.5, P value<0.0001), while competition without leachate reduced dry weight to a level between the two leachate treatments. Interestingly, plant height was not significantly reduced plant height, albeit plants in all treatments were shorter than the control (Fig 3 B). Leaf area was significantly reduced by irrigating with leachate (Fig 3 C) (F=10.4., P value<0.0001). Competition without leachate resulted in a reduction between the two leachate treatments (Fig 3 D). Ambient stomatal conductance (F=5.4, P value=0.0023) and PSII activity (not significant) were reduced only when comparing the leachate + competition treatment to the control, with both the leachate and competition treatments in between the two (Fig 3 D, E). Notably, the competition and leachate were mostly similar, and the combination of both did not result in a cumulative decrease, perhaps pointing to a saturation of neighbor response. Between the two lines selected for this experiment, PI564163 (BTx623) produced plants with significantly higher shoot and root dry weight, leaf area, and height (Fig S2). However, no significant interaction between genotype and competition/leachate was found.

**Figure 3:**
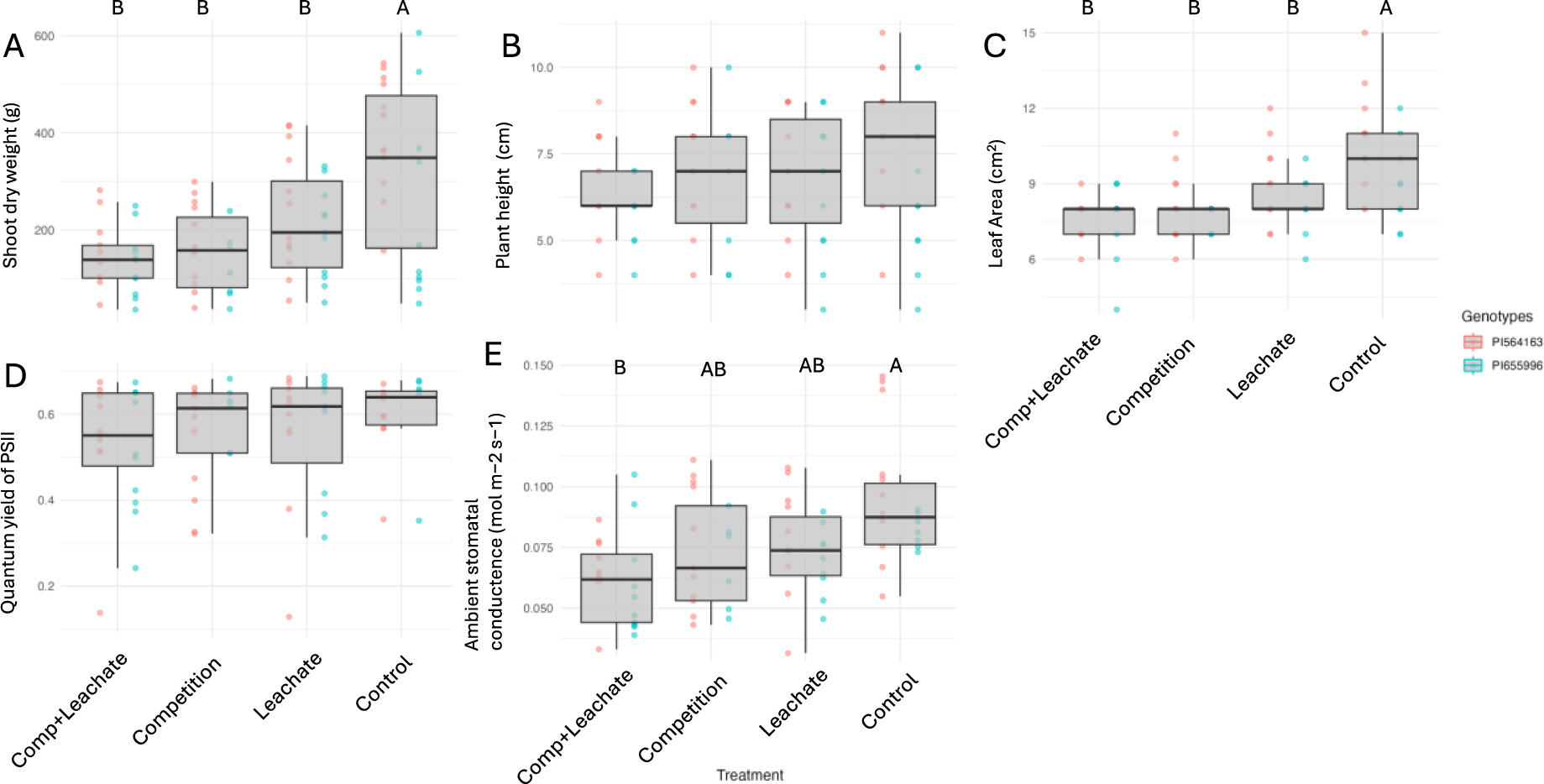
Shoot dry weight (A), plant height (B), leaf area (C), quantum yield of photosystem II, and Ambient stomatal conductance (E) of sorghum plants of two genotypes grown as a single plant (control) or a plant surrounded by neighbors (competition) and irrigated with a nutrient solution passed through a pot full of media alone (control and competition) or pots with sorghum plants (leachate-including root exudates, and leachate + competition). Different letters represent a significant difference between treatments in a Tukey HSD test (P<0.05).

Steady-state photosynthesis measurements were taken to test the changes in photosynthetic ability under different treatments (Fig 4). Our measurements show that assimilation rate (F=12.34.4, P value<0.0001), stomatal conductance (F=11.1, P value=0.0002), and transpiration (F=12.1, P value<0.0001) rate were all significantly reduced under the leachate treatment compared to the control and further significantly lowered under leachate + competition, with the competition (no leachate) treatment in between the two (Fig4 A, B, and D). Instantaneous water use efficiency was unaffected by any treatments, suggesting a reduction in assimilation was proportional to a reduction in stomatal conductance (Fig 4 C).

**Figure 4:**
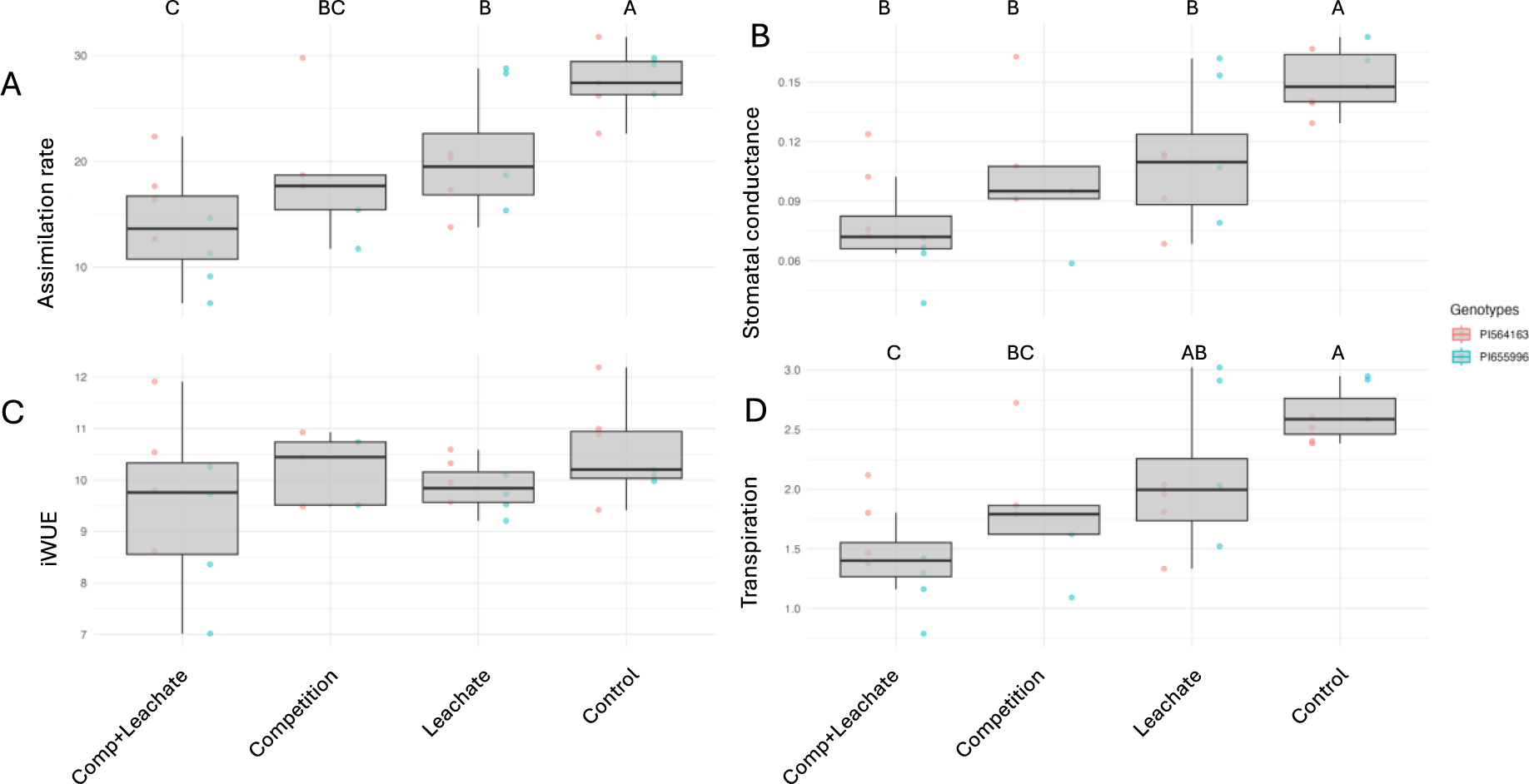
Steady-state assimilation rate (µmol CO2 m-2 s-1) (A), stomatal conductance (mol m-2 s-1)(B), instantaneous water use efficiency (iWUE, µmol CO2 m-2 s-1 /mmol H2O m-2 s-1)(C), and transpiration rate (mmol m-2 s-1) (D) of sorghum plants of two genotypes grown as a single plant or a plant surrounded by neighbors and irrigated with a nutrient solution passed through a pot full of media (control) or pots with sorghum plants growing in the same media (leachate-including root exudates). Different letters represent a significant difference between treatments in Tukey HSD tests (P<0.05).

We found a trend that competition reduced root diameter, root tip number, and total root length, but this effect was not statistically significant (Fig 5). The results in all measured root traits were consistent, albeit not statistically significant. Root traits showed high variability among individuals, suggesting a greater sample size would be required to conclusively demonstrate the effects of neighbors and leachate on root architecture.

**Figure 5:**
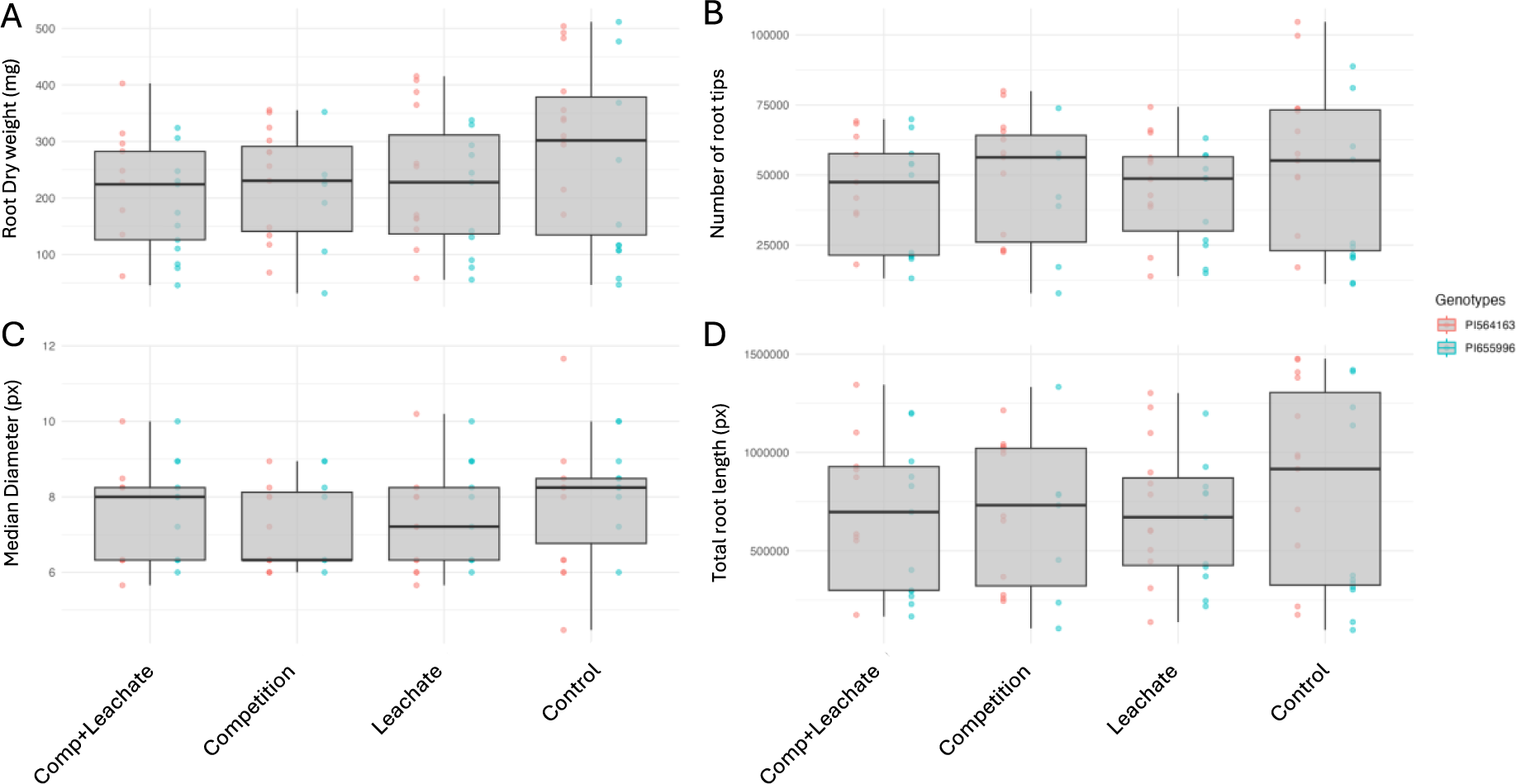
Root biomass in grams (A), number of root tips in pixels (B), median root diameter in pixels (C), and total root length in pixels (D) of sorghum plants of two genotypes grown as a single plant or a plant surrounded by neighbors and irrigated with a nutrient solution passed through a pot full of media (control) or pots with sorghum plants growing in the same media (leachate-including root exudates). Root architecture analysis was conducted using the RhizoVision Explorer v2.0.3 software.

## Discussion

This study sought to investigate the belowground effect of the presence of neighbors without nutrient deficiency in combination with changes in water availability. We tested whether root sensing of neighbors could change plant growth and productivity when there are ample resources. Further, we tested whether this signal could be carried in leachates likely containing rhizodeposits, without the physical presence of neighboring roots.

In the physical interaction experiment (EXP1), competition reduced sorghum productivity by similar levels to drought (Fig 1, Table 1), despite both water and nutrients being provided in ample supply, and light competition was suppressed by tying back neighbors. This indicates that focal plants responded to the presence of neighbors in the absence of soil resource depletion in a way that reduced productivity. This should be considered along with reported reductions in productivity under increased densities (more neighbors) for maize (Sarlangue *et al*. 2007) and sorghum (Jahanzad *et al*. 2013), to provide a new avenue for genotype evaluation.

To further study the nature of responses found in EXP1, the effect of competition was divided into signaling and physical presence. In the leachate transfer experiment (EXP2), source plants were irrigated with a nutrient solution every two days and showed no signs of stress. The leachate transfer experiment was designed to focus on the below-ground communication between plants, which is an important communication channel (Bais *et al*. 2004; Khashi U Rahman *et al*. 2019). This method also allowed us to study the root system of plants grown with and without the presence of signals from neighboring plants’ roots. The productivity and photosynthetic activity of plants exposed to competitors or leachates were lower in above-ground metrics (Fig 3,4); the similarity in response suggests root signaling. Such root-root signaling could potentially occur via cytokinin sensing (Bollmark and Eliasson 1986), which is present in root exudates and uptaken by roots (Soejima *et al*. 1992; Zahir *et al*. 2001). Further, the movement of cytokinin from root to shoot can cause changes in stomatal conductance and productivity (Glanz-Idan *et al*. 2020). In our study, the presence of leachate or competitors reduced below-ground biomass and led to changes in root architecture features, however, below the significance threshold (Fig 5).

The similarity in response between treatments with physical neighboring plants, in EXP1 and EXP2, and irrigation with leachate indicates the existence of a root-based biochemical response independent of biotic limitations or stress (Fig 2, 3). Other species, including grasses, have shown belowground biochemical signaling can reduce plant productivity. For example, A study of rice (*Oryza sativa*) showed that plants produced fewer tillers in response to neighbors producing active strigolactones in the growth media and that mutants in the D14 receptor of SL did not respond to neighbors (Yoneyama *et al*. 2022). Several mechanisms that should be considered in the context of plant-plant communication below ground include stress signals transferred from the source to the target pots. Another possible pathway for plant-plant communication, which leachates would carry, is the transfer of microbes and their metabolites. All pots were filled with the same new media, and the plant-plant interaction occurred within the same species. Thus, we do not believe the decreased performance is an effect of the microbes’ transfer between pots. However, through the feedback of microbes with other microbes, including changes to quorum sensing, microbes may be a vector for plants sensing neighbors.

The above-ground nature of the response to neighbors (reduced stomatal conductance, reduced size and leaf area), suggests an avoidance response when a plant senses neighbors, perhaps limiting their photosynthetic activity and reducing resource demand, will, in turn, require smaller root systems. This has been shown to contribute to facilitation in multispecies communities (Postma and Lynch 2012). While this could have a negative effect on an individual plant level, this type of strategy may support increased field-level or population productivity and represent a different aspect of increased fitness that might have been selected for in a crop (Lynch 2013). A possible benefit for the individual plant could arise from limiting their development in response to neighbors and thus avoiding local resource depletion. Selecting for smaller plants can be observed when considering studies pointing to higher productivity under increased densities in sorghum (Habyarimana *et al*. 2004) and maize (Cox 1996).

The importance of increasing the productivity of field crops, for which higher plant densities have been a common strategy since the middle of the last century (Donald 1968), is still a target for modern breeders. With the exception of (Lynch 2013) those who focused on root architecture and anatomy and their importance in an ideotype for high-density growing, most investigations of crop traits have focused on above-ground phenotypes. Thus, while above-ground interactions between plants have been studied (through shading), less is known about below-ground interactions between plants. Further, most ideotypes target stand-level productivity and do not focus on individual plants and their responses to the environment. Root system development in densely planted crops must support two sets of requirements: (a) uptake of necessary water and nutrients and (b) low interference with neighboring crop plants. These roles have led to a call for ideotypes of a root system that is highly branched at shallow depth but penetrates deep into the soil (Lynch 2013); such a root system would absorb nutrients in the plant’s immediate environment but not compete with neighbors, potentially increasing stand-level production (Lynch 2013; Mi *et al*. 2016).

The results reported here, obtained in systems that allowed for plant-plant interaction to occur below ground, suggest that efforts should be put into examining the response to neighbors regarding root development, which may hold a large potential for crop improvement. The variation in response to below-ground sensing neighbors found here (Fig 2) represents a possible tool for studying these responses to identify possible targets for breeding plants better suited for extremely high densities. If breeders follow our findings and test for below-ground interactions, they could inform their selection for higher-yielding lines through a pathway that does not change the commonly harvested parts of the plant.

## Acknowledgments

The authors thank Scott DiLoreto and his team for the technical support of this experiment, Jesica Janakiraman, and Rohan Ramaswamy for their assistance with sampling and plant care. Shiran Ben-Zeev was supported by BARD, the United States—Israel Binational Agricultural Research and Development Fund, Vaadia-BARD postdoctoral fellowship award number FI-625-2022.

## Supplementary Information

**Table S1:** Sorghum landraces used in the experiment.

| **Line code** | **PI** | **Name** | **Origin** | **Remarks from USDA grin/ sorghum collection (where applicable)** |
| --- | --- | --- | --- | --- |
| **L1** | PI 655996 | RTx430 | Texas | vegetative drought tolerance, natural LGS1 LOF |
| **L2** | PI565121 | Macia | Zimbabwe | background of LGS1 mutants |
| **L3** | PI564163 | BTx623 | Texas | Pre-flowering drought tolerant |
| **L4** | PI656025 | Shanqui Red | China | high striga germination |
| **L5** | PI533810 | Karad 2-7-11 | India | Plants short-statured, Photoperiod insensitive |
| **L6** | PI576364 | Chari Uri | India | Fertility reaction Maintainer |
| **L7** | PI656027 | SRN-39 | Sudan | low striga germination |
| **L8** | PI533752 | SC103 | South Africa | low striga germination, short-statured, Photoperiod insensitive |
| **L9** | PI533769 | 290 Feterita Shendi 2 | Sudan | Plants short-statured, Photoperiod insensitive. |
| **L10** | PI534070 | BE 25 | Nigeria | Plants short-statured, Photoperiod insensitive. |

**Table S2:**
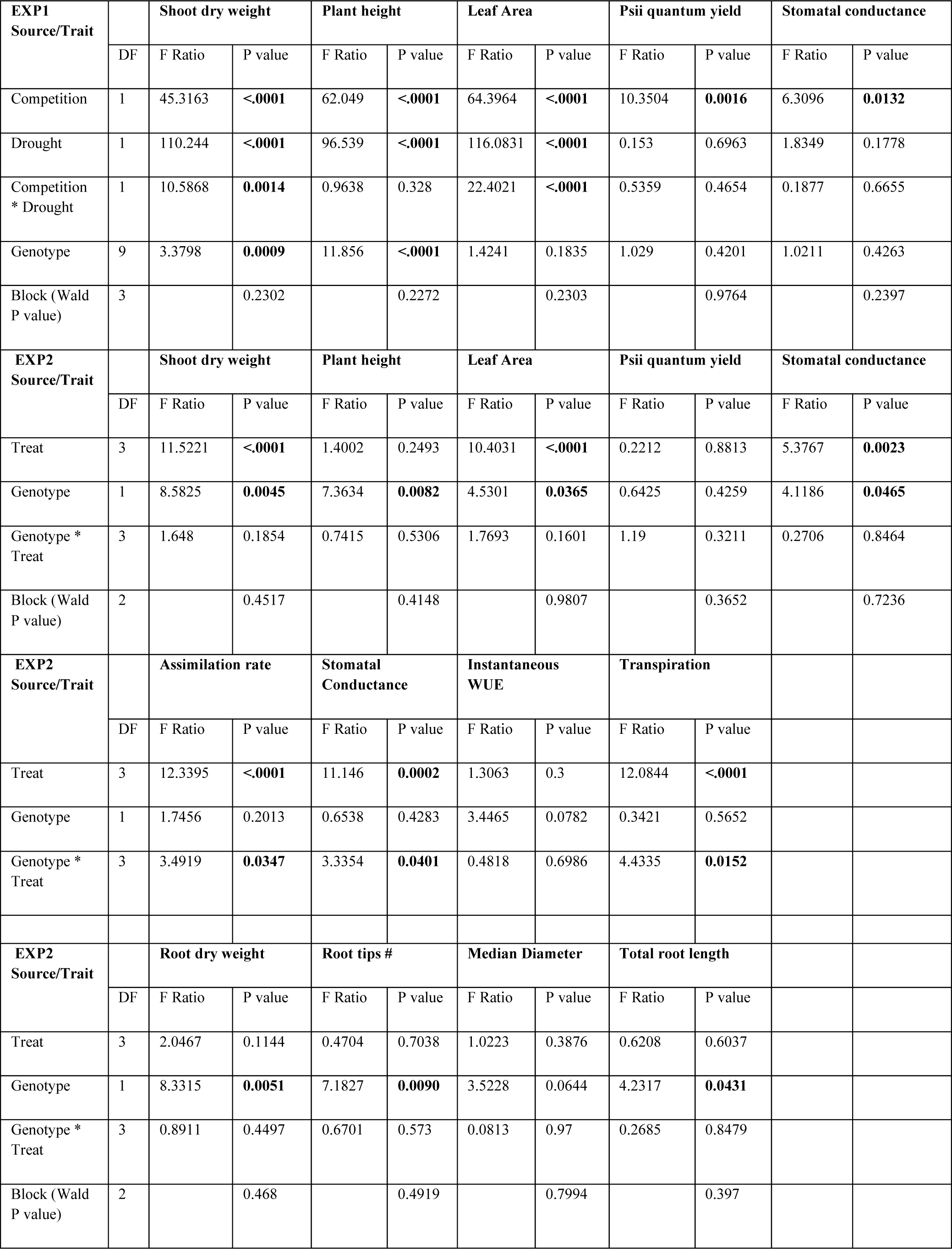
Sources of variance and significance in Experiment 1 (top section) and 2(bottom three sections ANOVA results. Genotype and treatment main effects are presented along with the Tukey HSD significance test (F ratio) and P values.

| EXP1<br>Source/Trait |  | Shoot dry weight |  | Plant height |  | Leaf Area |  | Psii quantum yield |  | Stomatal conductance |  |
| --- | --- | --- | --- | --- | --- | --- | --- | --- | --- | --- | --- |
|  | DF | F Ratio | P value | F Ratio | P value | F Ratio | P value | F Ratio | P value | F Ratio | P value |
| Competition | 1 | 45.3163 | <.0001 | 62.049 | <.0001 | 64.3964 | <.0001 | 10.3504 | **0.0016** | 6.3096 | **0.0132** |
| Drought | 1 | 110.244 | <.0001 | 96.539 | <.0001 | 116.0831 | <.0001 | 0.153 | 0.6963 | 1.8349 | 0.1778 |
| Competition * Drought | 1 | 10.5868 | **0.0014** | 0.9638 | 0.328 | 22.4021 | <.0001 | 0.5359 | 0.4654 | 0.1877 | 0.6655 |
| Genotype | 9 | 3.3798 | **0.0009** | 11.856 | <.0001 | 1.4241 | 0.1835 | 1.029 | 0.4201 | 1.0211 | 0.4263 |
| Block (Wald P value) | 3 |  | 0.2302 |  | 0.2272 |  | 0.2303 |  | 0.9764 |  | 0.2397 |
| EXP2<br>Source/Trait |  | Shoot dry weight |  | Plant height |  | Leaf Area |  | Psii quantum yield |  | Stomatal conductance |  |
|  | DF | F Ratio | P value | F Ratio | P value | F Ratio | P value | F Ratio | P value | F Ratio | P value |
| Treat | 3 | 11.5221 | <.0001 | 1.4002 | 0.2493 | 10.4031 | <.0001 | 0.2212 | 0.8813 | 5.3767 | **0.0023** |
| Genotype | 1 | 8.5825 | **0.0045** | 7.3634 | **0.0082** | 4.5301 | **0.0365** | 0.6425 | 0.4259 | 4.1186 | **0.0465** |
| Genotype * Treat | 3 | 1.648 | 0.1854 | 0.7415 | 0.5306 | 1.7693 | 0.1601 | 1.19 | 0.3211 | 0.2706 | 0.8464 |
| Block (Wald P value) | 2 |  | 0.4517 |  | 0.4148 |  | 0.9807 |  | 0.3652 |  | 0.7236 |
| EXP2<br>Source/Trait |  | Assimilation rate |  | Stomatal Conductance |  | Instantaneous WUE |  | Transpiration |  |  |  |
|  | DF | F Ratio | P value | F Ratio | P value | F Ratio | P value | F Ratio | P value |  |  |
| Treat | 3 | 12.3395 | <.0001 | 11.146 | **0.0002** | 1.3063 | 0.3 | 12.0844 | <.0001 |  |  |
| Genotype | 1 | 1.7456 | 0.2013 | 0.6538 | 0.4283 | 3.4465 | 0.0782 | 0.3421 | 0.5652 |  |  |
| Genotype * Treat | 3 | 3.4919 | **0.0347** | 3.3354 | **0.0401** | 0.4818 | 0.6986 | 4.4335 | **0.0152** |  |  |
| EXP2<br>Source/Trait |  | Root dry weight |  | Root tips # |  | Median Diameter |  | Total root length |  |  |  |
|  | DF | F Ratio | P value | F Ratio | P value | F Ratio | P value | F Ratio | P value |  |  |
| Treat | 3 | 2.0467 | 0.1144 | 0.4704 | 0.7038 | 1.0223 | 0.3876 | 0.6208 | 0.6037 |  |  |
| Genotype | 1 | 8.3315 | **0.0051** | 7.1827 | **0.0090** | 3.5228 | 0.0644 | 4.2317 | **0.0431** |  |  |
| Genotype * Treat | 3 | 0.8911 | 0.4497 | 0.6701 | 0.573 | 0.0813 | 0.97 | 0.2685 | 0.8479 |  |  |
| Block (Wald P value) | 2 |  | 0.468 |  | 0.4919 |  | 0.7994 |  | 0.397 |  |  |

**Figure S1:**
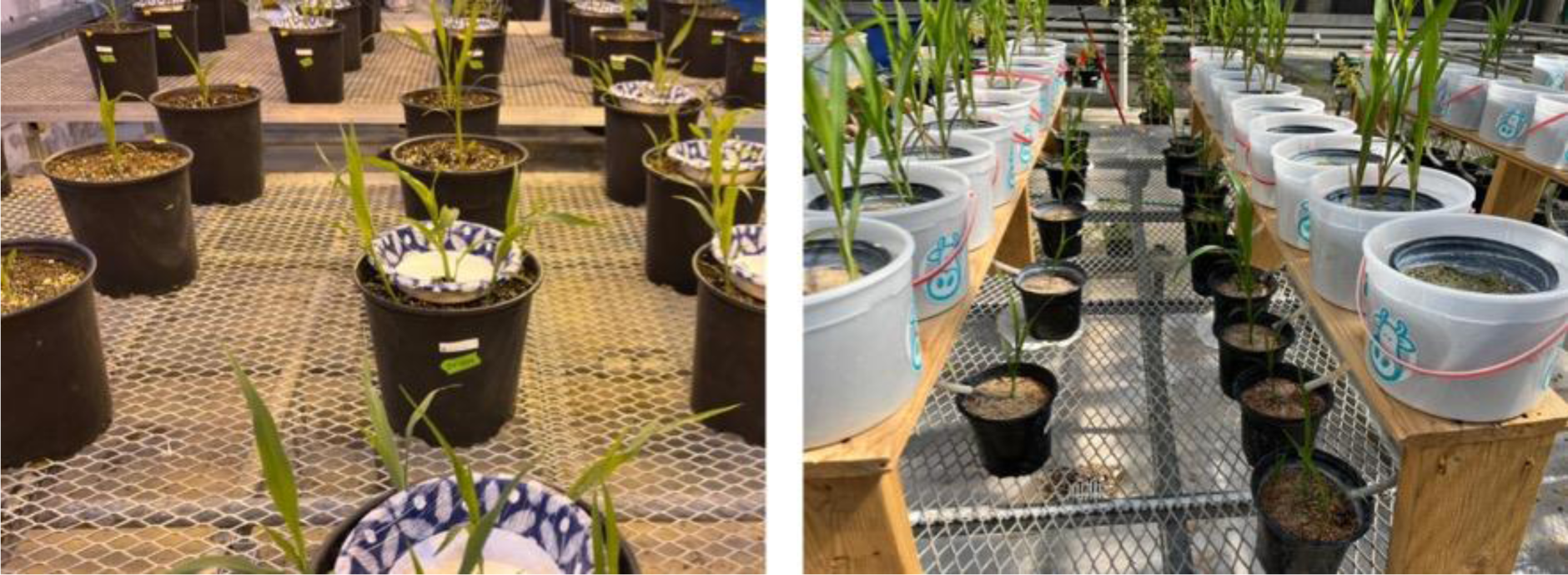
an overview of the experimental setup used for the physical competition experiment (left) and leachate transfer (right).

**Figure S2:**
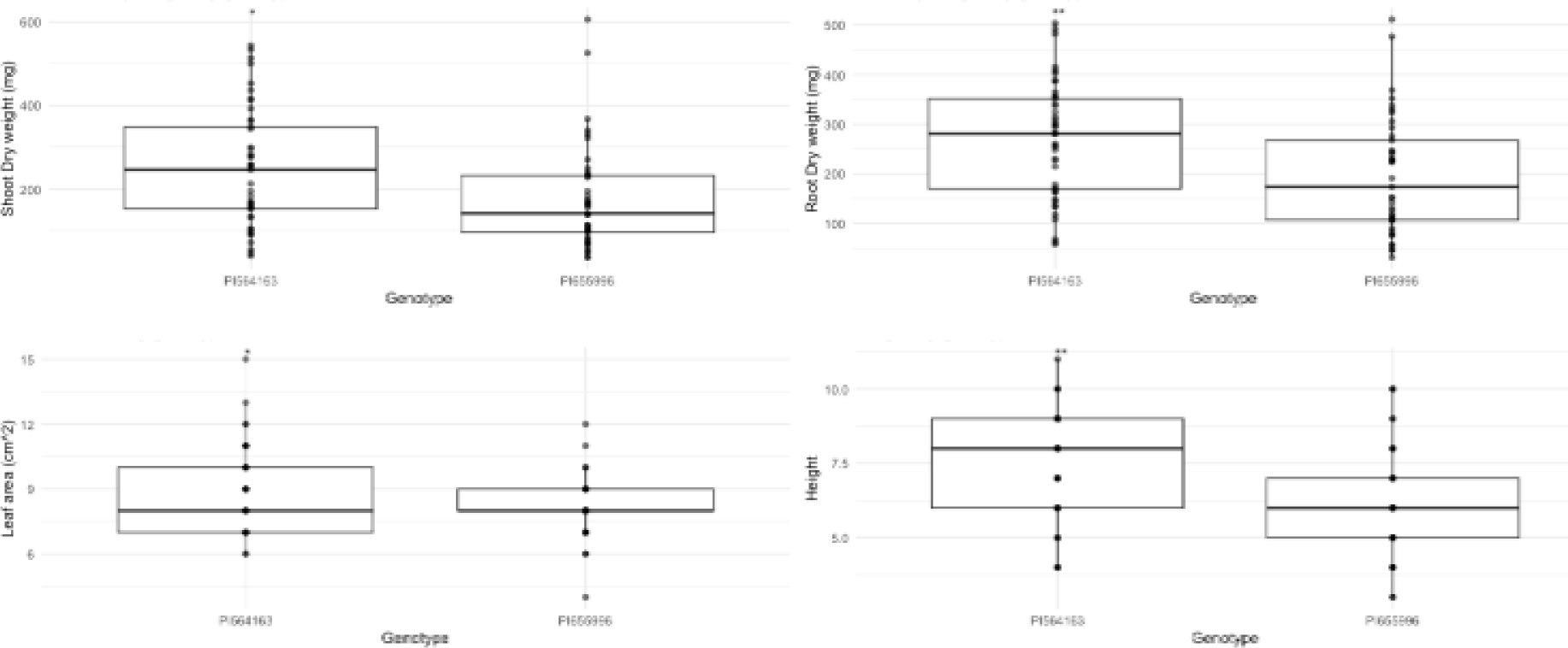
Shoot (top left) and Root (top right) dry weight, Leaf area (bottom left), and plant height (bottom right) of the two sorghum genotypes tested in the leachate transfer experiment (EXP2) - BTx623 and RTx430 (PI 564163 and 655996, respectively). “*”, “**”, and “***” indicate t-test P values <0.05, <0.01, and <0.001, respectively.

